# OGT (*O*-GlcNAc Transferase) selectively modifies multiple residues unique to lamin A

**DOI:** 10.1101/206458

**Authors:** Dan N. Simon, Amanda Wriston, Qiong Fan, Jeffrey Shabanowitz, Alyssa Florwick, Tejas Dharmaraj, Sherket B. Peterson, Yosef Gruenbaum, Cathrine R. Carlson, Line M. Grønning-Wang, Donald F Hunt, Katherine L. Wilson

**Affiliations:** Department of Cell Biology, Johns Hopkins University School of Medicine, Baltimore MD 21205 USA; Department of Chemistry, University of Virginia, Charlottesville VA 22904 USA; Department of Nutrition, Institute of Basic Medical Sciences, University of Oslo, 0317 Oslo Norway; Department of Biological Chemistry, Johns Hopkins University School of Medicine, Baltimore MD 21205 USA; Department of Genetics, Institute of Life Sciences, Hebrew University of Jerusalem, Givat Ram Jerusalem 91904 Israel; Institute for Experimental Medical Research, Oslo University Hospital and University of Oslo, 0450 Oslo, Norway; Department of Pathology, University of Virginia, Charlottesville VA 22904 USA

**Keywords:** Lamin, Nuclear lamina, *O*-GlcNAcylation, *O*-linked N-acetylglucosamine (*O*-GlcNAc) Transferase (OGT)

## Abstract

The *LMNA* gene encodes lamins A and C with key roles in nuclear structure, signaling, chromatin organization, and genome integrity. Mutations in *LMNA* cause >12 diseases, termed laminopathies. Lamins A and C are identical for their first 566 residues. However, they form distinct filaments *in vivo* with apparently distinct roles. We report that lamin A is *O*-GlcNAc modified in human hepatoma (Huh7) cells and in mouse liver. In vitro assays with purified OGT enzyme showed robust *O*-GlcNAcylation of recombinant mature lamin A tails (residues 385-646), with no detectable modification of lamin B1, lamin C, or ‘progerin’ (Δ50) tails. Using mass spectrometry, we identified 11 *O*-GlcNAc sites in a ‘sweet spot’ unique to lamin A, with up to seven sugars per peptide. Most sites were unpredicted by current algorithms. Double-mutant (S612A/T643A) lamin A tails were still robustly *O*-GlcNAc-modified at seven sites. By contrast, *O*-GlcNAcylation was undetectable on tails bearing deletion Δ50, which causes Hutchinson-Gilford progeria syndrome, and greatly reduced by deletion Δ35, suggesting this region is required for substrate recognition or modification by OGT in vitro. These results suggest OGT, an essential protein and master regulator, regulates partners or function(s) unique to lamin A that are lost in progeria.

## INTRODUCTION

Lamins, encoded by three human genes (*LMNA*, *LMNB1*, *LMNB2*), form nuclear intermediate filaments that support nuclear structure, cell mechanics, development, genome organization, DNA repair, signaling, and tissue-specific gene silencing (1–3). Lamins have a conserved molecular structure; each polypeptide has a small globular ‘head’ domain, a long coiled-coil ‘rod’ domain, and a ‘tail’ comprising an Ig-fold domain as well as in the case of lamin A, an extended unstructured region (4–6). Mutations in *LMNA* cause diverse tissue-specific diseases (7). Some ‘laminopathies' affect mainly striated muscle (e.g., Emery-Dreifuss muscular dystrophy; dilated cardiomyopathy) whereas others perturb metabolism, causing insulin resistance syndrome (8) or Dunnigan-type familial partial lipodystrophy (FPLD2). FPLD2 is a puberty-onset disorder characterized by lipodystrophy, muscle hypertrophy, and insulin-resistant diabetes (9,10), and elevated (dysregulated) hepatic glucose production (11). *LMNA* missense mutations are also reported in patients with metabolic syndrome (12,13), although genetic causality has not been established. In rare cases, *LMNA* mutations cause Hutchinson-Gilford Progeria Syndrome (HGPS) or related phenotypes (14). Most HGPS patients have a mutation that alters pre-mRNA splicing, generating a 50-residue deletion (‘Δ50’) lacking the site required for ZMPSTE24-dependent proteolytic maturation of the lamin A precursor (15,16). The resulting permanently-farnesylated protein, named ‘progerin’, has acute and long-term effects on nuclear structure and function (3,17). About 35% of progerin-expressing HGPS patients are also insulin resistant (18).

These metabolic phenotypes drew our attention to evidence that lamin A is modified by a nutrient stress-responsive enzyme named OGT (*O*-GlcNAc transferase), which adds a simple sugar, *O*-GlcNAc (β-*O*-linked N-acetylglucosamine), to Serine or Threonine residues of target proteins. Endogenous lamin A is *O*-GlcNAc modified at two sites (S612 and T643) in mitotic HeLa cells (19). *O*-GlcNAc is a ubiquitous, reversible, and dynamic modification that controls signaling, transcription, and mitosis (20). *O*-GlcNAc is added to target proteins by OGT, and removed by *O*-GlcNAcase (OGA), each encoded by a single essential gene in mammals (21–23). Notably, OGT is ‘tuned’ to metabolism in liver, kidney, and pancreatic-cells; its catalytic activity increases in tandem with cellular levels of glucose or glucosamine, which are converted via the hexosamine biosynthetic pathway to UDP-GlcNAc, the donor substrate for *O*-GlcNAcylation reactions (24–26). OGT activity can protectively increase during stress, including recovery of the heart from ischemic injury (27). However, long term cardiomyopathy associated with diabetes, aging, and hypertension all correlate with aberrantly high levels of protein *O*-GlcNAcylation, suggesting OGT hyper-activity perturbs heart function (28). Protein hyper-*O*-GlcNAcylation has pathophysiological implications in insulin resistance, non-alcohol fatty liver disease, fibrosis, and is also characteristic of diabetic nephropathy (29), the leading cause of chronic kidney disease and a significant long-term complication of diabetes.

We investigated potential *O*-GlcNAc modification of A-type lamins in hepatoma (Huh7) cells and liver tissue. We also tested recombinant purified tail domains of A-and B-type lamins as potential substrates for OGT *in vitro*. We report extensive *O*-GlcNAc modification of residues unique to lamin A *in vitro*, and evidence that lamin A substrate recognition by OGT requires residues deleted in HGPS.

## RESULTS

To assess potential *O*-GlcNAcylation of A-type lamins, we cultured human hepatoma (Huh7) cells for 24 h in medium containing physiological (5 mM) or high (25 mM) glucose. A-type lamins were immunoprecipitated from sonicated Huh7 nuclear lysates using monoclonal antibody 5G4, which recognizes the Ig-fold domain (residues 466-484; Supplemental Figure 1A) in both lamin A and lamin C. Immunoprecipitates (50% per lane) were resolved by SDS-PAGE with input lysate controls (8%), and immunoblotted (Figure 1A). One set of samples was probed using antibody CTD_110.6_, which recognizes the *O*-GlcNAc modification, then stripped and reprobed for lamins A/C using antibody 5G4 (Figure 1A). The ~73 kD lamin A band was consistently *O*-GlcNAc-positive in both concentrations of glucose (asterisk; Figure 1A; n= 5), demonstrating *O*-GlcNAc modification of lamin A in cultured human hepatoma cells. However, the level of lamin A *O*-GlcNAcylation in 5-vs-25 mM glucose, and the extent to which other *O*-GlcNAcylated proteins co-immunoprecipitated with lamin A in 25 mM glucose, varied between experiments and was not studied further. *O*-GlcNAc signals were not detected on lamin C (~60 kD; Figure 1A). These results demonstrated *O*-GlcNAcylation of endogenous lamin A in proliferating hepatoma cells, consistent with previous results from mitotic HeLa cells (19).

**Figure 1.**
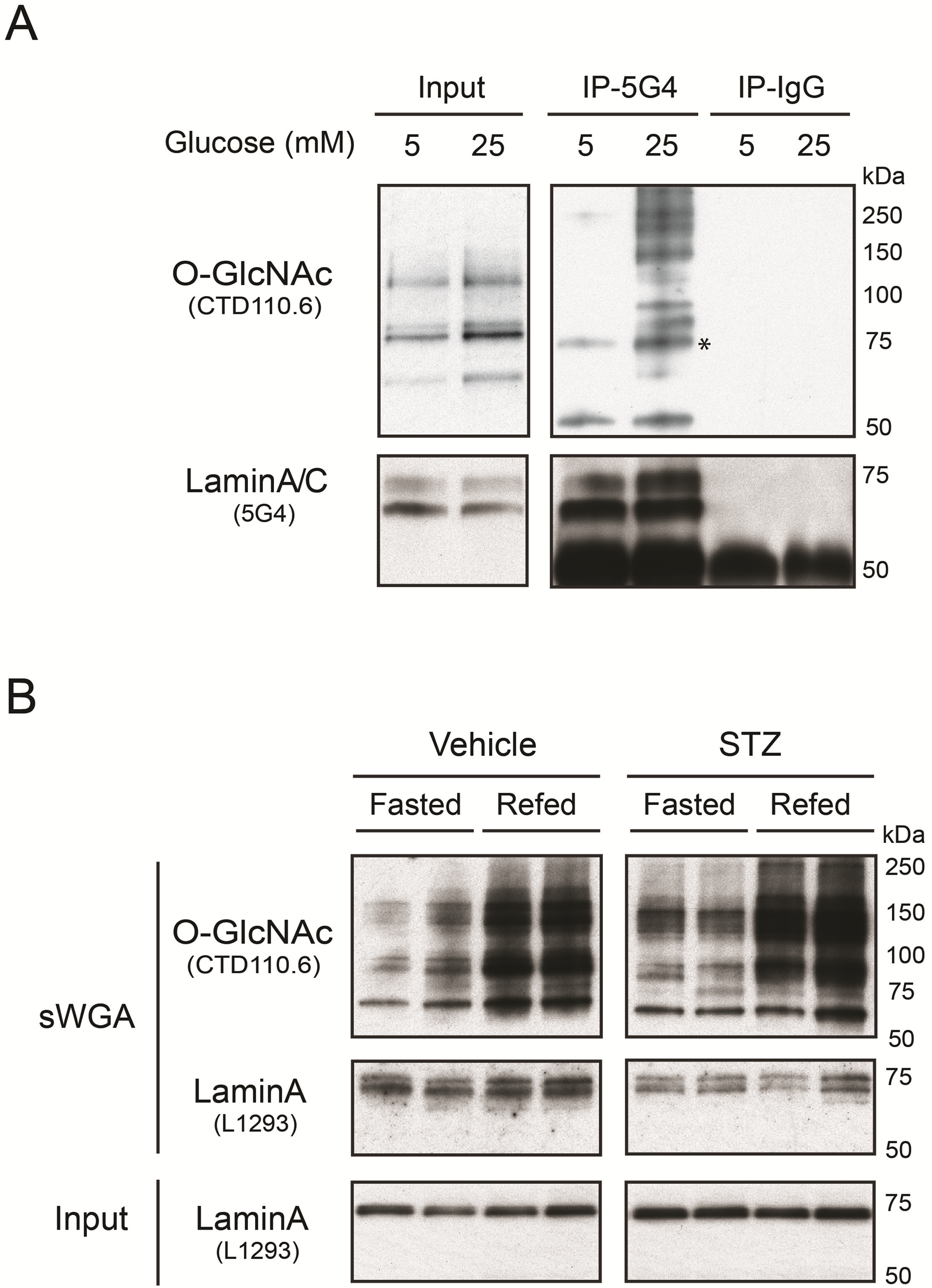
*O*-GlcNAcylation of native lamin A in human hepatoma (Huh7) cells and mouse liver. (**A**) Western blots of proteins from human hepatoma (Huh7) cells, cultured 24 h in physiological (5 mM) or high (25 mM) glucose, probed with antibody CTD_110.6_ specific for the *O*-GlcNAc modification (*O*-GlcNAc), then stripped and reprobed with antibody 5G4, specific for lamins A and C (LaminA/C). Control input lysates, and proteins immunoprecipitated using lamin A/C antibody 5G4 or nonspecific IgG as the negative control, were resolved in parallel gels. Asterisk indicates the *O*-GlcNAc signal corresponding to immunoprecipitated lamin A. (**B**) Western blots of mouse liver nuclear proteins, before (Input) or after affinity purification on sWGA (sWGA), which binds *O*-GlcNAc-modified proteins. We analyzed two livers for each condition. Male mice were either fasted (Fasted), or fasted and then refed for 12 h (‘Refed’), and were either pretreated with streptozotocin to induce hyperglycemia (‘STZ’), or not (‘Vehicle’). Input controls were probed with lamin A-specific antibody L1293. Affinity-purified samples were probed first for the *O*-GlcNAc modification (antibody CTD_110.6_), then stripped and reprobed for lamin A (antibody L1293). Control input lysates, and proteins immunoprecipitated using lamin A/C antibody 5G4 or nonspecific IgG as the negative control, were resolved in parallel gels.

To determine whether lamin A was *O*-GlcNAcylated in a non-proliferating tissue, we analyzed liver lysates from wildtype mice that were either fasted for 24 h, or fasted 24 h and then re-fed for 12 h on a regular chow diet to boost overall protein *O*-GlcNAcylation (see Methods). *O*-GlcNAc-modified proteins from each nuclear lysate (two livers per condition) were selectively affinity-purified using succinylated Wheat Germ Agglutinin (‘sWGA’; does not bind sialylated proteins). The resulting *O*-GlcNAc-modified proteins were then resolved by SDS-PAGE and western blotted for *O*-GlcNAc (antibody CTD_110.6_) and lamin A (antibody L1293; Figure 1B). Control input lysates were analyzed in parallel. Lysates contained similar levels of input mature lamin A (~73 kD; Figure 1B, Input). As expected, re-feeding caused an overall increase in *O*-GlcNAcylation of nuclear proteins (CTD_110.6_ Ab; Figure 1B). Probing for specific proteins in the affinity-purified samples revealed enrichment for two lamin A-specific bands: the expected ~73 kD mature lamin A band and a slightly larger (~74 kD) band (sWGA; Figure 1B). This ~74 kD band, which might represent prelamin A or a posttranslationally-modified form of mature lamin A, was not further characterized. We found similar results (sWGA purification of two presumably *O*-GlcNAc-modified forms of lamin A) in liver lysates from mice treated with streptozocin (STZ; Figure 1B), which kills pancreatic Δ-cells and models diabetes (30). The relative abundance of the two lamin A bands could not be interpreted, because the core epitope recognized by antibody L1293 (residues 604-611; Supplemental Figure 1) adjoins a potential *O*-GlcNAc site (S612; (19)) and potential phosphorylation sites (31,32), and may be differentially accessible on native lamin A. Collectively these results from hepatoma cells and mouse livers support the hypothesis that lamin A is *O*-GlcNAc-modified, and hence *O*-GlcNAc-regulated, in the liver. To explore whether OGT targets lamin A selectively, we compared the tail domains of A-and B-type lamins as biochemical substrates for OGT *in vitro*.

### *In vitro O*-GlcNAcylation of recombinant lamin tails

To identify *O*-GlcNAc sites, we incubated recombinant lamin tail domains with purified OGT *in vitro*. We tested nine different tail constructs: lamin B1, lamin C, mature lamin A residues 385-646 (wildtype, S612A, T643A, or S612A/T643A), and prelamin A residues 385-664 bearing the progeria-associated Δ35, Δ50, or Δ90 deletions (Figure 2A). Each purified recombinant T7-tagged lamin tail polypeptide (1 μg protein) was incubated with UDP-GlcNAc and calf intestinal phosphatase (CIP), plus or minus purified recombinant His-tagged OGT enzyme, for 2 h at 22-24°C, then overnight at 4°C. Reactions were quenched with SDS sample buffer, resolved by SDS-PAGE and immunoblotted first with antibody CTD_110.6_, specific for the *O*-GlcNAc moiety, then stripped and reprobed with antibodies against the T7-tag on lamins. These *in vitro O*-GlcNAcylation assays revealed robust and specific modification of the recombinant mature lamin A tail domain (Figure 2B). *O*-GlcNAcylation of mature lamin A tails mutated at one or both previously-reported *O*-GlcNAc sites (S612A, T643A, S612A/T643A) was similarly robust (Figure 2B), suggesting OGT targeted alternative *O*-GlcNAc sites. The *O*-GlcNAc signals were specific: potential antibody cross-recognition of recombinant proteins was ruled out by probing duplicate membranes with the *O*-GlcNAc antibody plus 100 mM free sugar (+GlcNAc; Figure 2C). *In vitro O*-GlcNAcylation was greatly reduced by deletion Δ35 (loss of residues 622-658; (16)): trace signals were detected only in long exposures (Figure 2B; ‘long exp’). There was no detectable *O*-GlcNAcylation of the HGPS-associated deletion Δ50 (Δ608-658; (15)) or deletion Δ90 (Restrictive Dermopathy-associated loss of residues 567-658; (33)) *in vitro* (Figure 2B,C; n=3). There were also no specific *O*-GlcNAc signals on lamin C or lamin B1 tails (Figure 2B), or on full-length *C. elegans* lamin (‘ce-lamin’; Figure 2D), which has a short ‘B-like’ tail. Potential *O*-GlcNAc modification of the head or rod domain of human lamins was not tested. These results did not rule out potential *O*-GlcNAc modifications of lamin C or B-type tails, since negative results can be artifacts of recombinant proteins. However, selective *O*-GlcNAcylation of lamin A tails *in vitro* was consistent with our results in hepatoma cells (Figure 1A). These results suggested OGT targets the tail domain of lamin A selectively *in vitro*, and is sensitive to progeria-associated deletions.

**Figure 2.**
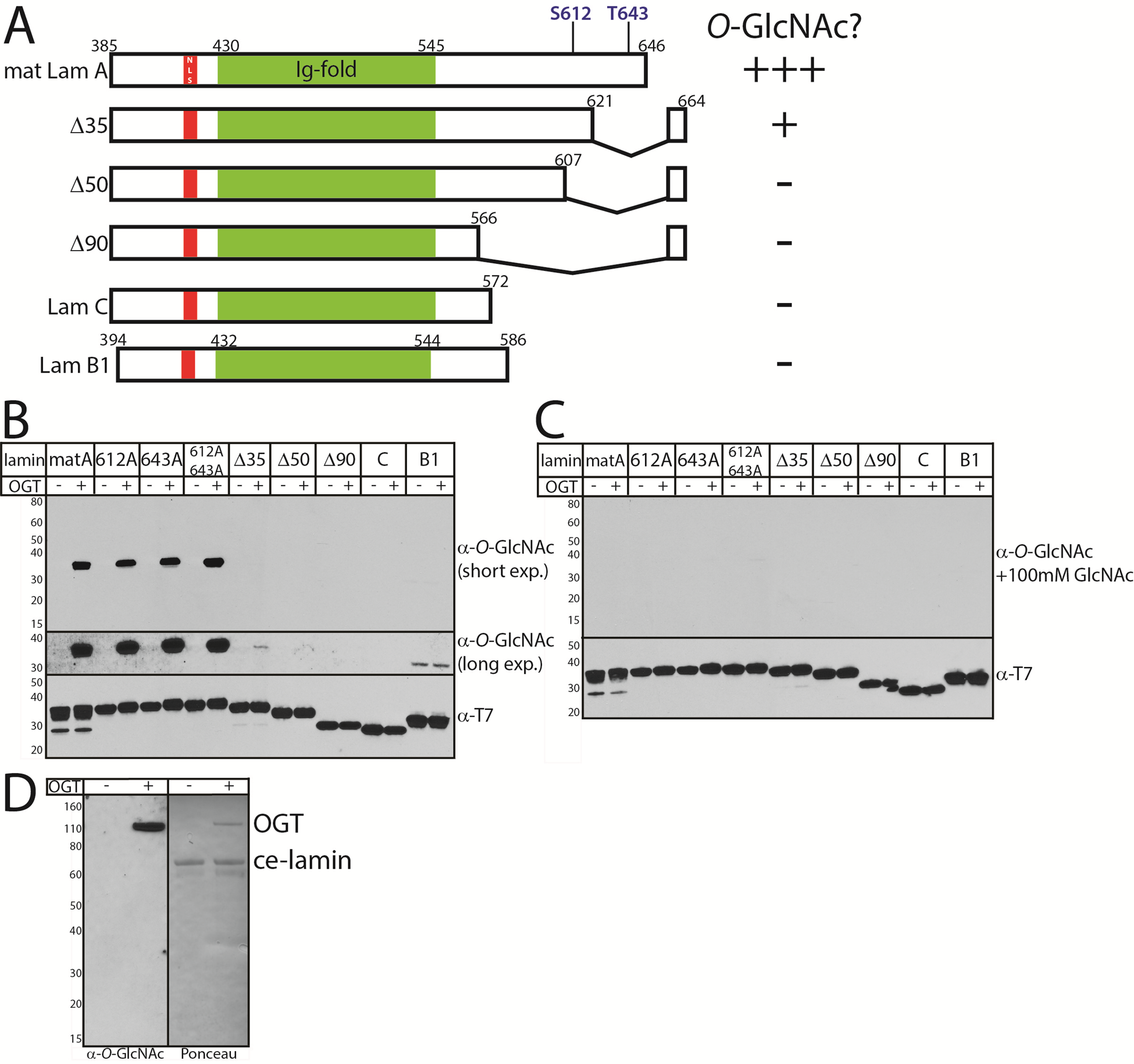
*In vitro* O-GlcNAcylation of recombinant lamin tails. **(A)** Schematic diagram of the recombinant lamin tail constructs used as substrates for *in vitro*
modification by recombinant OGT enzyme. These included mature wildtype lamin A tail residues 385-646 (‘mat LamA’); mature lamin A residues 385-646 bearing single or double Ala-substitutions at S612 and T643 (19); deletions Δ35, Δ50, or Δ90 in the context of prelamin A tail residues 385-664; and wildtype lamin C (‘lam C’) and lamin B1 (‘lam B1’) tails. The nuclear localization signal (NLS) and Ig-fold domain (92) are indicated. Results for each construct are summarized at right. **(B, C)** *In vitro O*-GlcNAcylation results, assayed by western blotting with the CTD_110.6_ *O*-GlcNAc antibody. Lamin tails (wildtype, S612A, T643A, S612A/T643A, Δ35, Δ50, Δ90; lamin C or B1) were incubated with UDP-GlcNAc and CIP with or without OGT for 2 h (22-25°C), then overnight at 4°C. Reactions were resolved by SDS-PAGE and immunoblotted with antibody CTD_110.6_ (α-*O*-GlcNAc) in the absence (**B**) or presence of competing 100 mM free sugar (**C**), then stripped and re-probed with T7-tag antibodies (α-T7-tag) to detect lamin tails. **(D)** *In vitro O*-GlcNAcylation results for recombinant full-length *C. elegans* lamin, detected by Ponceau-staining prior to immunoblotting for *O*-GlcNAc, were negative. OGT activity was confirmed by auto-*O*-GlcNAcylation of the OGT band.

### *O*-GlcNAc site identification by mass spectrometry

*O*-GlcNAc sites were identified by mass spectrometry analysis of three *in vitro*-modified samples: wildtype mature lamin A tail, S612A/T643A-mutated mature lamin A tail, and pre-lamin A bearing the Δ35 HGPS-causing deletion. Samples were digested with either AspN or AspN followed by chymotrypsin. The generated peptides were analyzed by on-line HPLC-MS/MS on a high-resolution mass spectrometer. Peptides were fragmented by both collisionally activated dissociation (CAD) and electron transfer dissociation (ETD) in a data-dependent mode. We achieved nearly 90% peptide coverage of both the wildtype and S612A/T643A-mutated lamin A tails. Coverage of Δ35 was lower (71%) but included 48 of 57 total Ser/Thr residues in the Δ35 tail (Figure 3A). The *O*-GlcNAcylated peptides were identified from CAD spectra by either the O-GlcNAc oxonium signature peak at m/z 203 amu, or they exhibited the charge reduced product ion with the loss of 203 amu, as shown in Figure 4A. ETD spectra were used to identify sites modified by *O*-GlcNAc. We detected a total of 11 unique *O*-GlcNAc sites in the wildtype lamin A tail, four of which were more abundant, based qualitatively on the relative abundance of the ions corresponding to the *O*-GlcAc sites: S612, S613, S618, and T643 (‘more abundant’; Figure 3B). The seven less abundant sites were S603, S619, T623 and four others (S615, S616, T621, S628) and were detected only on heavily *O*-GlcNAc-modified peptides (‘less abundant’; Figure 3B). These results validated previously reported sites S612 and T643 (19) and identified nine more *O*-GlcNAc sites in the mature lamin A tail.

**Figure 3.**
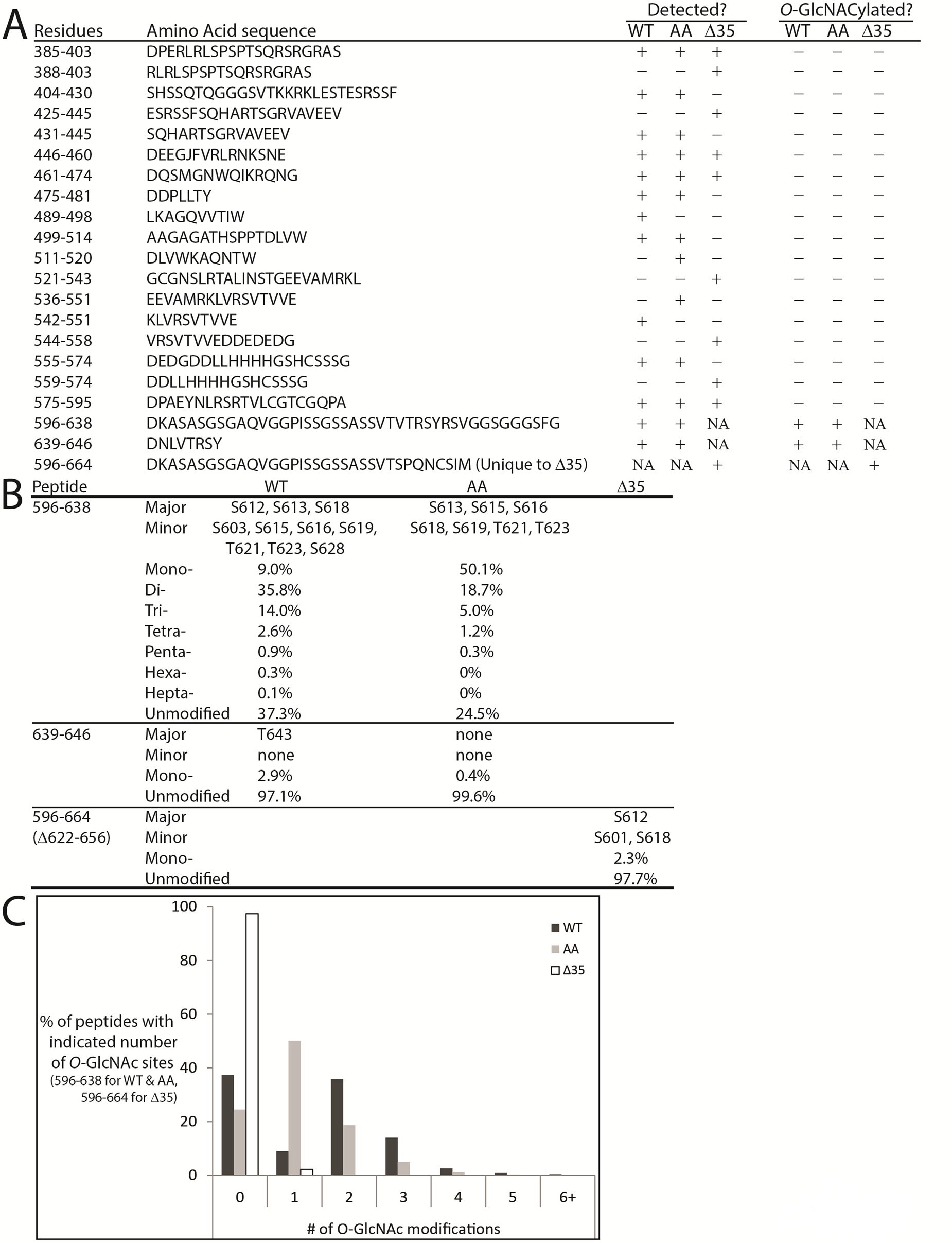
Mass spectrometry identification of *O*-GlcNAc sites in recombinant lamin A tail polypeptides. (**A**) Combined sequence coverage of AspN and AspN-Chymotrypsin digests of lamin A tails. Note: peptide DNLVTRSY (residues 639-646) included a vector-derived C-terminal residue L647. (**B**) Summary and relative abundances of identified *O*-GlcNAc sites in each indicated peptide from wildtype, double-mutant (S612A/T643A), or Δ35 lamin A tails. (**C**) For peptides comprising wildtype lamin A residues 596-638 or the corresponding double-mutant (S612A/T643A) peptide 596-638, the percentages of peptides with zero (unmodified), one, two, or more *O*-GlcNAc-modified sites are graphed.

Lamin A tail peptides were frequently modified at multiple sites (up to seven; Figure 3B,C). Mass spectrometry analysis of the AspN peptide containing residues 596-638 from wildtype lamin A revealed that 37.3% were unmodified, 9% had one *O*-GlcNAc, 35.8% were di-GlcNAcylated, 14% were tri-GlcNAcylated, and the remaining 3.9% had four to seven *O*-GlcNAc modifications (Figure 3B,C). Percentages are based on the chromatographic peak area of each species compared to the chromatographic peak area of all species combined. Only two di-GlcNAcylated pairs were detected: S612+S613 and S612+S603. The ETD spectrum of the S612+S613 di-GlcNAcylated pair, with near complete sequence coverage, is shown in Figure 4B. The most frequent triple combinations were S612+S613 plus one of S616, T623, or S603; together these five sites accounted for all triples in the wildtype tail.

**Figure 4.**
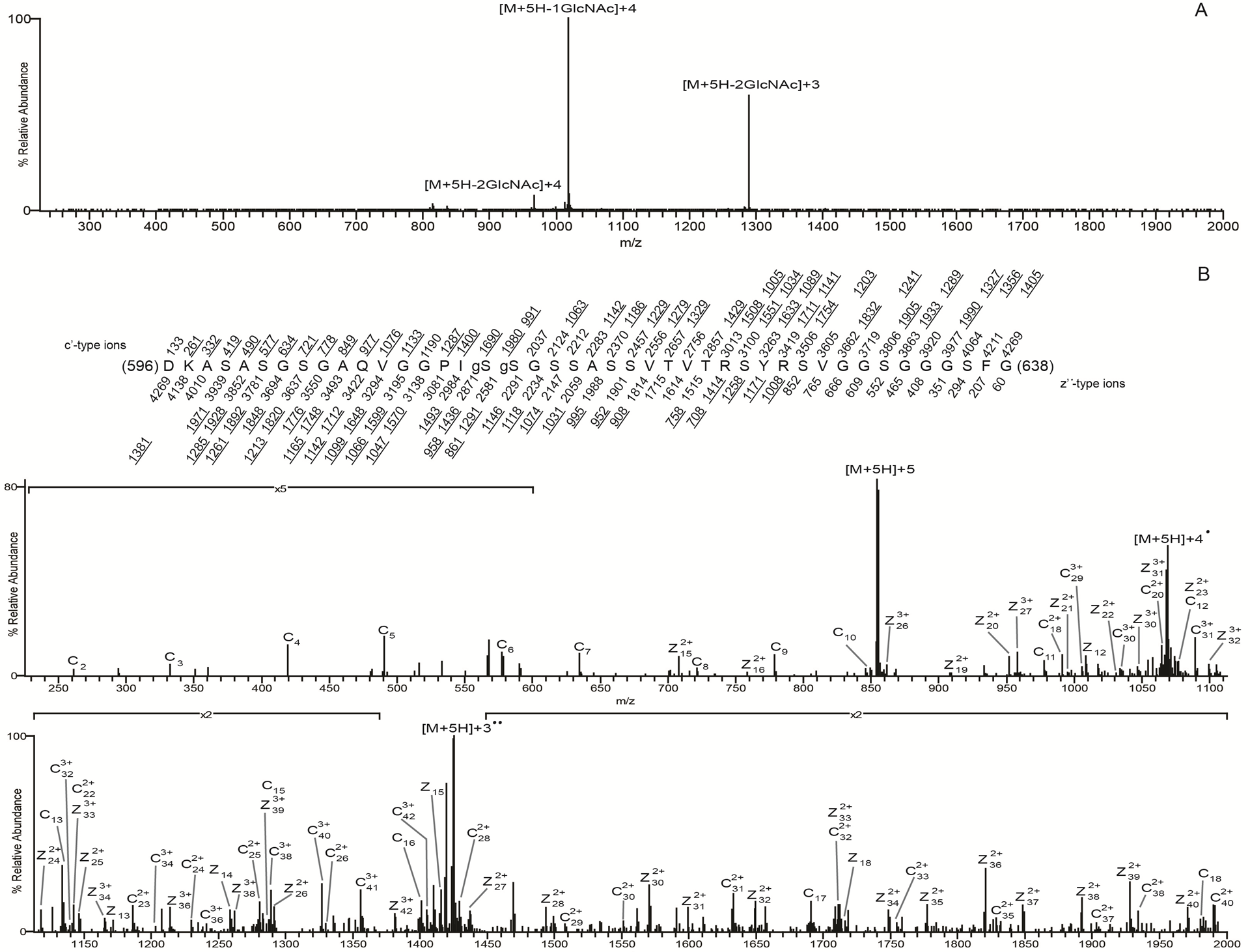
*O*-GlcNAc site mapping on the wildtype lamin A tail via liquid chromatography-MS/MS. (**A**) CAD MS/MS spectrum recorded on the (M+5H)^5+^ ions (m/z 855.2046) of the di-GlcNAcylated peptide DKASASGSGAQVGGPIgSgSGSSASSVTVTRSYRSVGGSGGGFG. The CAD spectrum contains the charge-reduced ions minus the loss of 1GlcNAc, 203 Da, at m/z 1018 for the +4 species, and the charge-reduced ion minus the loss of 2GlcNAc residues, 406 Da, at m/z 1289 for the +3 species and 967 for the +4 species. (**B**) An ETD MS/MS spectrum recorded on the [M+5H]^5+^ ions (m/z 855.2046) of the di-GlcNAcylated peptide DKASASGSGAQVGGPIgSgSGSSASSVTVTRSYRSVGGSGGGFG. Predicted product ions of types c’ - and z’^⋅^ - are listed above and below the peptide sequence, respectively. Singly charged ions are listed as monoisotopic masses and doubly and triply charged ions are listed as average masses. ETD product ions are labeled in the ETD spectrum. Observed product ions are underlined and are sufficient to define the O-GlcNAc residues at Ser612 and Ser613, indicated by ‘g’ in the figure.

We also identified sites in the corresponding AspN peptide from the double-mutant (S612A/T643A; ‘AA’) tail, to determine how OGT responded to substrates lacking both previously identified *O*-GlcNAc sites (19). The mono-GlcNAc species (invariably at S613) predominated (50.1%) over the unmodified species (24.5%; Figure 3B). The di-and tri-GlcNAc species were also relatively abundant (18.7% and 5%), and tetra-and penta-GlcNAc species were present at <2% each (‘AA’; Figure 3B). Collectively >25% of double-mutant tails had two or more modifications (Figure 3B). Mass spectrometry analysis of the double mutant revealed *O*-GlcNAc modifications at three predominant sites (S613, S615, S616) and four less abundant sites (S618, S619, T621, T623; Figure 3B). We were unable to site-map the mono-*O*-GlcNAcylated peptide corresponding to residues 639-646 due to poor sequence coverage; however, given the T643A mutation, this peptide had only one potential site, S645, suggesting *O*-GlcNAc-modification of residue S645 in the double-mutant. Modification at S645 was not detected in wildtype or Δ35 tails. We concluded that the S612A/T643A double-mutant tail remained attractive as a substrate for OGT.

With the S612A/T643A-mutated substrate, OGT apparently compensated by targeting three predominant sites adjoining residue 612 (S613, S615, S616) plus nearby residues S618, S619, T621, and T623. However, there were fewer ‘multiples’: the peptide comprising residues 596-638 was more likely to have one modification (50.1% versus 9% of wildtype), and less likely to have two (18.7% vs 35.8% of wildtype), three (5% vs 14% of wildtype), or four or more modifications (1.5% vs 3.9% of wildtype; Figure 3B). In the peptide comprising residues 596-626, we detected di-GlcNAcylation at residues S613+S616 and S613+S615, as well as tri-GlcNAcylation at S613+S616+S618 and S613+S615+T621. In a longer version of this peptide (residues 596-638), we detected di-GlcNAcylation mainly on S613+S615 and less often on S613+(S616 or S618 or S619). Figure 5A shows the CAD spectrum of the di-GlcNAcylated species with the most abundant peaks being the charge reduced product ion with the loss of 203 and 406, one and two O-GlcNAc moieties, respectively. Figure 5B shows the ETD spectrum of the S613 and S618 di-GlcNacylated peptide with almost full sequence coverage. Tri-GlcNAcylation occurred on S613 plus different combinations of S615, S616, T621, and/or T623 as the second and third sites. We speculate that the first *O*-GlcNAc modification (e.g., S612 in wildtype; S613 in the AA mutant) strongly favors further modification.

**Figure 5.**
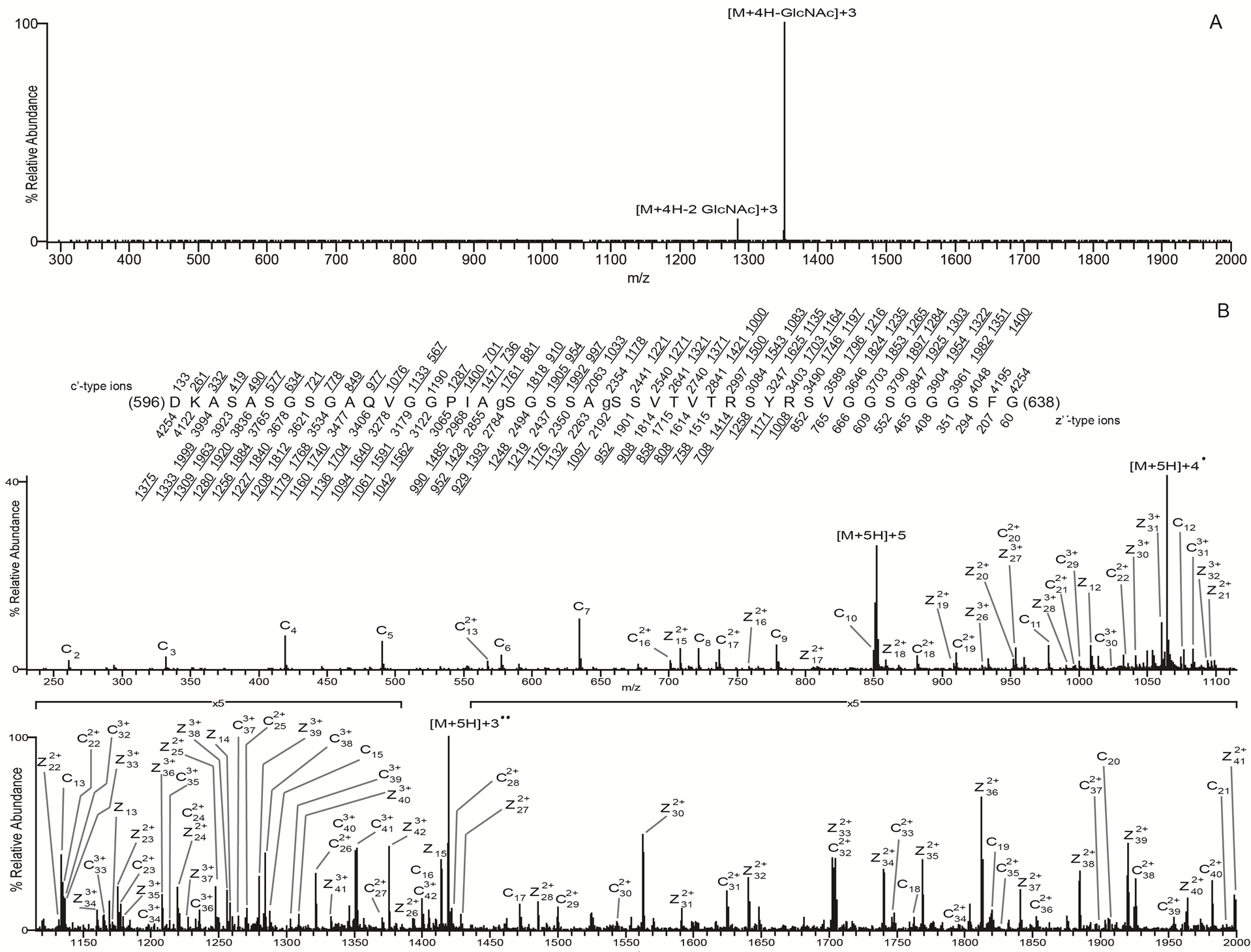
*O-*GlcNAc site mapping on the S612A/T643A-mutated mature lamin A tail via liquid chromatography-MS/MS. (**A**) CAD MS/MS spectrum recorded on the (M+4H)^4+^ ions (m/z 1064.2561) of the di-GlcNAcylated peptide DKASASGSGAQVGGPIAgSGSSAgSSVTVTRSYRSVGGSGGGFG. The CAD spectrum contains the charge-reduced ions minus the loss of 1GlcNAc, 203 Da, at m/z 1351, and the charge-reduced ion minus the loss of 2GlcNAc residues, 406 Da, at m/z 1284. (**B**) An ETD MS/MS spectrum recorded on the [M+5H]^5+^ ions (m/z 852.0084) of the di-GlcNAcylated peptide DKASASGSGAQVGGPIAgSGSSAgSSVTVTRSYRSVGGSGGGFG. Predicted product ions of types c’ - and z’^⋅^ - are listed above and below the peptide sequence, respectively. Singly charged ions are listed as monoisotopic masses; doubly and triply charged ions are listed as average masses. ETD product ions are labeled in the ETD spectrum. Observed product ions are underlined and are sufficient to define the *O*-GlcNAc residues at Ser612 and Ser618, indicated by ‘g’ in the figure.

The Δ35 tail, with nine hypothetical sites (see Figure 6), was modified very inefficiently: 97.7% of peptides were unmodified, with only single modifications detected at either S612 (<3% of peptide 596-664) or less frequently at S601 or S618 (Figure 3A,B). Peptides that included residue S613 were recovered, but unmodified, supporting the hypothesis that Δ35 greatly impairs substrate recognition or modification by OGT *in vitro*.

These results identify residues 601-645 as a broadly defined *O*-GlcNAc ‘sweet spot’ (Figure 6), since OGT targeted this region robustly even when two abundant sites (S612 and T643) were missing (‘AA’ mutant; Figures 2B and 3B). This sweet spot overlaps the region required for ZMPSTE24-dependent cleavage of prelamin A, raising the possibility that OGT might influence prelamin A maturation. However, the ‘sweet spot’ is also a permanent feature of mature lamin A (Figure 7), suggesting OGT regulates roles unique to lamin A.

**Figure 7.**
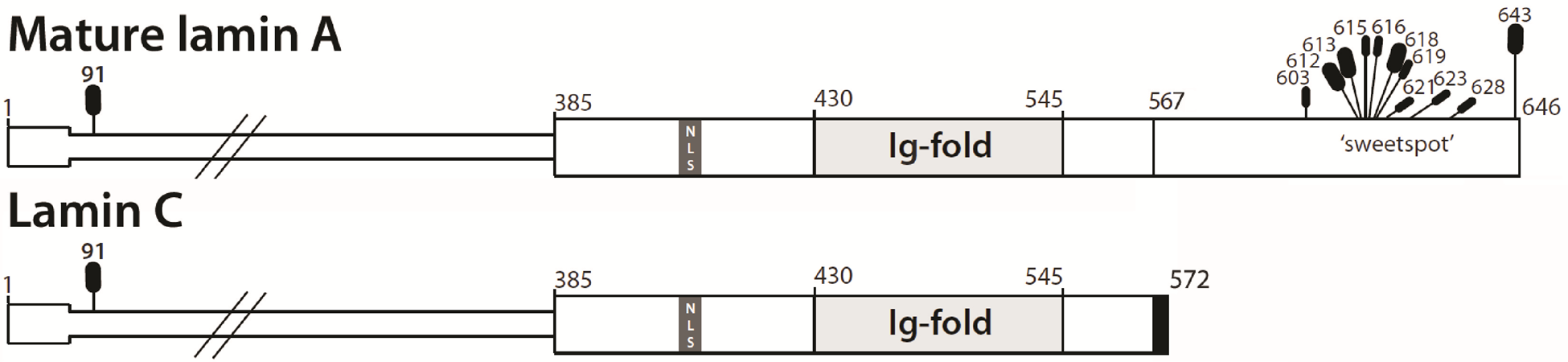
Schematic of the mature lamin A and lamin C proteins, which have identical residues 1-566, showing all currently identified *O*-GlcNAc sites. Long *vs* short black ovals indicate greater or lesser abundance at each identified site, determined experimentally for the lamin A tail (see text). The relative abundance at site T91, identified in embryonic stem cells (38), is not known. It is also not known if *O*-GlcNAcylation at T91 is restricted to one isoform (not depicted), or found on both proteins (as depicted here).

## DISCUSSION

Lamin glycosylation was first suggested nearly 30 years ago (34). The nature of this modification was unknown until *O*-GlcNAc sites were identified in mitotic HeLa cells (S612 and T643; (19)) and in mouse brain (sites S611 and S613, homologous to human S612; (35)). We found that endogenous lamin A is *O*-GlcNAcylated in human hepatoma cells and in mouse liver. Our biochemical analysis shows that OGT modifies lamin A tails selectively and robustly *in vitro*. Interestingly, all 13 identified primary and ‘compensatory’ *O*-GlcNAc sites are located within a ‘sweet spot’ (residues 601-645) unique to lamin A. This region includes ‘super-sweet’ residues 612-623, where *O*-GlcNAc was detected on every available Ser and Thr (Figure 6). Two of our four most abundant sites, S612 and T643, are biologically relevant (19,35). Our results independently confirm two previously-reported *in vitro* sites, S623 and S628 (36). We also identified seven novel *O*-GlcNAc sites in mature wildtype human lamin A (S603, S613, S615, S616, S618, S619, T621) and two novel compensatory sites: S601 (identified in Δ35) and S645 (identified in the S612A/T643A mutant). Our results do not rule out possible *O*-GlcNAc sites in the neck or Ig-fold domains, since we recovered the majority of - but not all - Ser/Thr-containing peptides from these regions. However, the lack of *O*-GlcNAcylation of either the 50, 90, or lamin C tails strongly implies that OGT selectively targets residues 601-645 in the lamin A tail. One limitation of this study is that we did not examine the head or rod domains, or precursor residues 647-664. These regions warrant further study, since residue T91, in the coil-1B domain shared by all A-type lamins (37), is *O*-GlcNAc-modified in human embryonic stem cells (38).

### Most *O*-GlcNAc sites in lamin A are not predicted by current algorithms

Substrate recognition by OGT is complex, involving both its TPR (protein-protein interaction) and catalytic domains, and can be influenced by post-translational modifications and partners (23,39). There is no strict consensus site for *O*-GlcNAc modification (39). For example, a study of pre-selected 13-mer peptides showed that *O*-GlcNAc sites are influenced by size and conformational restrictions in the −3 to +2 positions (36,40), and reported *O*-GlcNAc modifications at three sites (T623, S625, S628) in lamin A peptide ^618^SSVTV<u>**T**</u>R<u>**S**</u>YR<u>**S**</u>VG^630^ and one site (T399) in lamin B1 peptide ^389^KLSPSPSSRV<u>**T**</u>VS^401^ (36). Even though our lamin B1 tail polypeptide included T399, *O*-GlcNAcylation was not detected (Figure 2B), possibly because this fragment began at residue 394 and was potentially insufficient for OGT recognition. For lamin A, we identified T623 and S628 (but not S625) as low-abundance sites, in the context of our much longer (>250-residue) tail polypeptides. Overall, very few identified *O*-GlcNAc sites in lamin A are predicted by the Pathak et al. (2015) consensus. A shorter consensus proposed by Liu and colleagues (41) fared slightly better: most ‘sweet spot’ sites match at positions −1 and +2, but only three of our 13 sites match this consensus at position −2.

In agreement with Jochmann and colleagues (39), many sites on lamin A were not predicted. E.g., OGTsite (42) predicted three known sites (S612, T623, T643), and three sites we did not detect. The YinOYang *O*-GlcNAc prediction server (http://www.cbs.dtu.dk/services/YinOYang; 43) strongly predicted two identified sites (S612, T621), weakly predicted five identified sites (S603, S613, S615, S618, T623), failed to predict five sites (T91, S616, S619, S628, T643; 42% false negative rate) and predicted 8 other tail sites we did not detect. *O*-GlcNAcPRED (44) predicted only two sites (both known: S612, T643) at higher stringency; at lower stringency, this program predicted one more identified site (S601) plus three tail sites we did not detect. We suggest that *O*-GlcNAc sites identified in long polypeptides, rather than peptides, will be critical to improving predictive algorithms and understanding the molecular basis for OGT modifications at two or more adjoining residues, as seen for lamin A (this work) and its nuclear membrane partner, emerin (45).

### The ‘sweet spot’ in relation to the site of ZPMSTE24-dependent cleavage

Figure 6 shows the identified *O*-GlcNAc sites in relation to published constructs, expressed as GFP-fusions, that were cleaved by ZMPSTE24 with either 100% efficiency (‘41-mer’; residues 624-664), 50% efficiency (‘31-mer’; residues 634-664) or not cleaved (‘29-mer’; residues 636-664) in human embryonic kidney (HEK293) cells (46). The ‘super-sweet’ region directly borders the 41-mer region sufficient for ZMPSTE24-dependent cleavage (Figure 6, Prelamin A). Outside residues might also influence maturation, since a G608S substitution blocks FLAG-prelamin A cleavage in mouse embryonic fibroblasts (47). Speculatively, OGT access to prelamin A might be influenced by hypothetical OGT association with Zmpste24 (48; ‘PPI Finder’ algorithm). Whether OGT modifies prelamin A-specific residues 647-664, or influences lamin A maturation, are important open questions.

**Figure 6.**
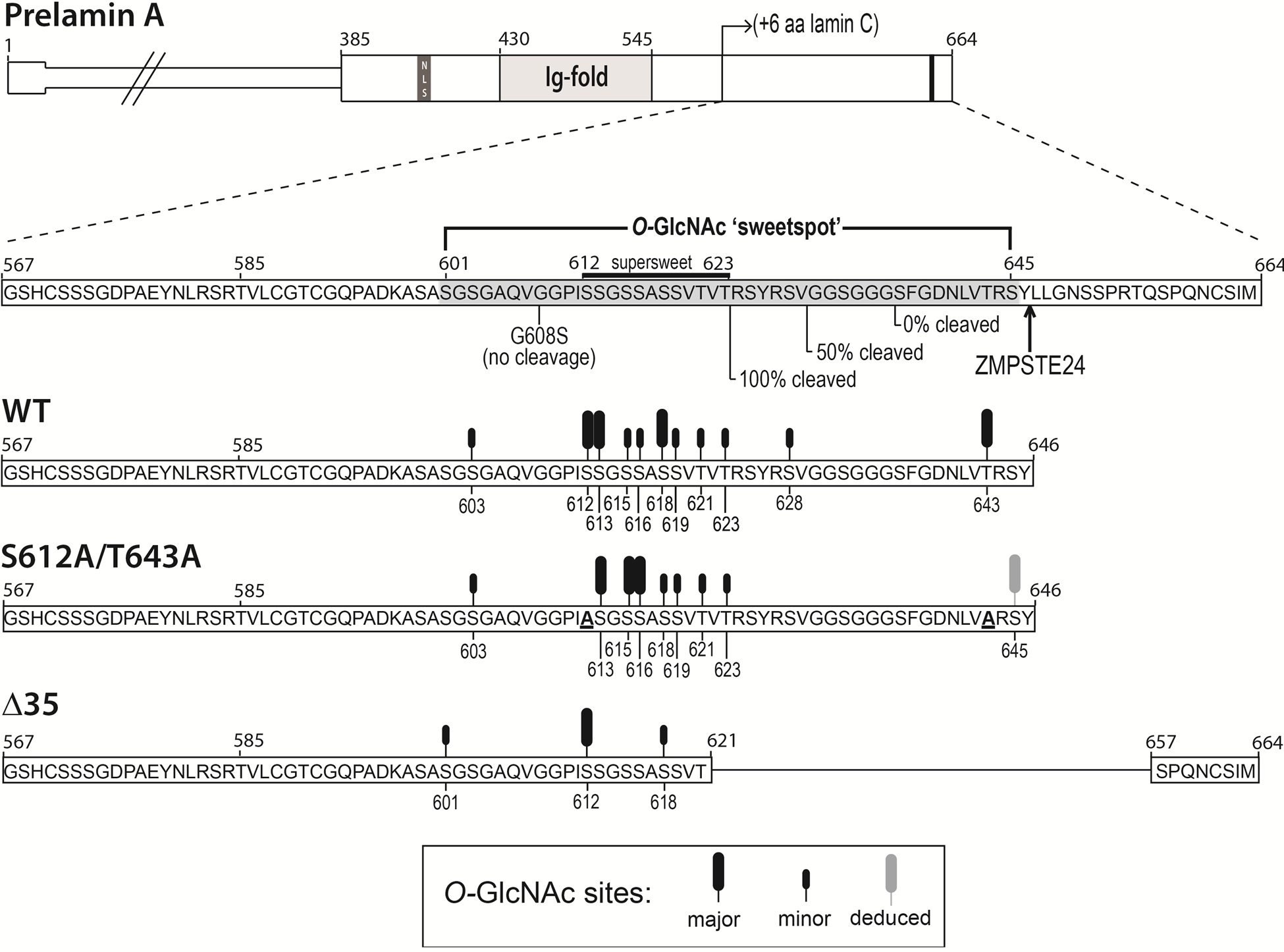
Schematic of lamin A tail polypeptides showing identified *O*-GlcNAc sites. (**A**) Schematic diagram of full-length prelamin A and the three recombinant lamin A tail fragments (wildtype [WT], S612A/T643A, Δ35) analyzed by mass spectrometry, all of which started with residue 385. All identified *O*-GlcNAc sites are located in the C-terminal region unique to prelamin A, expanded below (dotted lines) to show the amino acid sequence. Abundant or less-abundant *O*-GlcNAc sites are symbolized by large ('major’) or small ('minor’) black rods, respectively. The ZPMSTE24-dependent proteolytic cleavage site is indicated. The starting points of fragments that were cleaved by ZPMSTE24 with 100%, ~50% or 0% efficiency in cells (46) are indicated below the sequence. Mutation G608S is sufficient to block cleavage in a different assay (47). The region targeted by OGT in both wildtype and mutated lamin A tails (residues 601-645) is broadly designated the ‘sweet spot’, and includes ‘supersweet’ residues 612-623 (*O*-GlcNAc detected at every Ser and Thr).

### Selective OGT targeting of lamin A as a mechanism for separate functions and differential regulation of lamin A vs. lamin C

The discovery that lamin A and lamin C form separate filaments is so new and unexpected (49,50) that few studies have even attempted to distinguish one from the other. We found that OGT selectively modifies the tail domain of mature lamin A, but not lamin C, *in vitro*, raising the hypothesis that lamin A is uniquely and differentially regulated by OGT. These results add to growing evidence for functional specialization. For example, lamin C specifically associates with nuclear pore complexes (50). Specialization is also evident in the brain and in metabolism. Lamin C predominates in neurons, where mRNAs encoding prelamin A are eliminated by a specific microRNA (51). Compared to wildtype controls, mice that express lamin C as their only A-type lamin have significantly longer lifespans, despite accumulating white fat and developing abdominal tumors that limited their lifespan; these mice also have reduced rates of metabolism and respiration, fewer mitochondria, higher fasting insulin levels, and higher blood glucose levels (52). Mice that express mature lamin A-only have nuclear shape defects in fibroblasts, but otherwise normal body weight, health, and fertility (53), implying appropriate regulation of mature lamin A. A-type lamins are linked to the mTOR pathway, which senses glucose and other nutrients to promote biosynthetic pathways, and inhibit or limit catabolic (breakdown) pathways including autophagy (54–56). Autophagy is significantly elevated in *lmna*-mutated cells (57,58). Mice that express H222P-mutated *lmna* in the heart have elevated AKT and mTOR signaling that leads to cardiomyopathy and impaired fasting-induced autophagy (59). These phenotypes are ameliorated by mTOR inhibitors, suggesting an A-type lamin(s) normally dampens mTOR signaling (60–62). It will be interesting to determine how OGT, which responds independently to nutrient stress, influences lamin A under these conditions.

What happens if OGT cannot modify the lamin A tail? Two elements are lost by deletion in ‘classic’ (Δ50) HGPS: functions specifically mediated by the lamin A tail, and OGT-dependent control of these functions. Neither element is likely to be restored by therapies that block farnesylation. Treating HGPS patients with the farnesyltransferase inhibitor Lonafarnib improved weight gain (18) and reduced prevalence of stroke (63), but did not affect other parameters including insulin resistance (seen in 35% of patients (18)). We found that progeria-associated deletions either abolished (Δ50) or greatly reduced (Δ35) lamin A modification by OGT *in vitro*. Three missense variants that perturb metabolism – p.G602S (insulin resistance syndrome (8); type 2 diabetes (64)), p.G631D (metabolic syndrome (12)), and p.R644C (multiple phenotypes (65)) - are located in the ‘sweet spot.’ Variant p.L92F, identified in an obese patient with severe metabolic syndrome (13), adjoins the reported *O*-GlcNAc site at T91 (38). However more work is needed to decipher mechanisms, since variants that affect metabolism, like those that cause other laminopathies, can be located throughout the polypeptide.

### Potential impacts of *O*-GlcNAc on mature lamin A function

OGT and lamin A are each notable for fundamental and global roles that include signaling, epigenetic regulation, and mitosis (3,66,67). For example, lamins and key partners (LEM-domain proteins and BANF1) are required to dynamically rebuild nuclear structure during exit from mitosis (68–72). Similarly, OGT is required for dividing cells to survive (21,23). For lamins, specific roles appear to be timed and controlled by competing posttranslational modifications, the best understood of which is phosphorylation. From this perspective, we note that ten of our 12 total identified *O*-GlcNAc sites can be phosphorylated in a variety of cell types or conditions (31,32, www.phosphosite.org). To our knowledge, *O*-GlcNAc is the only reported modification of lamin A residues T91 (38) and T621 (this work). Our findings suggest that lamin A (like cytoplasmic intermediate filaments) is specifically and dynamically regulated by crosstalk between OGT and multiple kinases (73).

*O*-GlcNAc and phosphate modifications can profoundly alter the conformations of intrinsically disordered polypeptides, such as the lamin A tail (74,75), and hence dynamically regulate access by binding partners. Partners that bind the unstructured region of lamin A, besides ZPMSTE24 (discussed above), include SUN1 (76), Narf, RBBP4, PKCΔ, and actin (31). Certain partners identified in a high-throughput yeast 2 hybrid screen are disrupted by disease-associated missense variants in the ‘sweet spot’, and might therefore be influenced by *O*-GlcNAc modifications. These partners include LAP1β (disrupted by lamin A variant p.G602S), ZNF3 (disrupted by p.G602S, p.G608S, p.T623S, p.R644C), MORF4L1 (disrupted by p.G602S, p.G608S, p.T623S), Cyclin G1 (disrupted by p.G602S, p.G608S, p.R644C), and CENP-P (disrupted by p.R644C) (77). We predict ‘sweet-spot’ modifications are very likely to influence association with AIMP3, a scaffolding protein. Normally found in the cytoplasm with roles in translation, AIMP3 also enters the nucleus and acts as a tumor suppressor (79). AIMP3 binds directly to residues 641-647 in the ‘sweet spot’ of mature lamin A (78). AIMP3 recruits Siah1 (an E3 ubiquitin ligase) and triggers selective degradation of mature lamin A to promote senescence (78,79). Interestingly, *O*-GlcNAc modifications of the unstructured lamin A tail might also have a broader impact, by controlling its ability to ‘fold back’ and restrict access to binding sites on the Ig-fold domain, as suggested for actin (80) and the nuclear pore complex protein, Tpr (81).

OGT is a master regulator that responds directly to nutrient stress, with key roles in liver, kidney and pancreatic Δ-cells (82,83). Our discovery that lamin A is *O*-GlcNAc-modified in hepatoma cells and in liver raises the possibility, for individuals with wildtype *LMNA* genes, that lamin A becomes misregulated under conditions of metabolic stress, with potentially global and long-term effects on signaling, epigenetic regulation, and 3D chromatin organization (84-87).

## EXPERIMENTAL PROCEDURES

### Cell culture

Huh7 liver hepatoma cells grown at passages 15-30 were maintained in high glucose (25 mM) DMEM (D6546; Sigma-Aldrich) supplemented with 10% FBS, 4 mM L-glutamine, and 1% penicillin-streptomycin. Subconfluent cells in 10-cm culture dishes were adapted to physiological glucose (5 mM) DMEM (D6046; Sigma-Aldrich) overnight before culturing cells in physiological glucose or high glucose for 24 h.

### Nuclear extraction, immunoprecipitation, and western blotting

Nuclear lysates from Huh7 cells were prepared using the NE-PER extraction kit (Pierce Biotechnology) with the following inhibitors added: 1 mM NaF, 1 mM Na_3_VO_4_, 1 mM Δ-glycerophosphate, 1 uM GlcNAc-thiazoline (OGA inhibitor), and Complete^TM^ protease inhibitors (Roche Applied Science), followed by sonication on ice (30 sec on, 30 sec off per session; three sessions), and centrifugation (10 min, 16,000*g*). Supernatant (nuclear lysate) proteins were stored at −20°C. For each immunoprecipitation, 100 ug nuclear lysate proteins were pre-cleared by rotating with Protein A/G agarose (sc-2003; Santa Cruz Biotechnology) for 1 h (4°C), then centrifuged (2 min, 500*g*). Supernatants were incubated with mouse monoclonal antibody against human lamin A/C (clone 5G4, kindly provided by Robert Goldman, Northwestern University; 1 ug per reaction), rotated overnight (4°C), then added to 20 ul Dynabeads Protein G (Invitrogen) and rotated 1 h at 4°C. Immunoprecipitates were washed thrice in PBS with 0.2% NP-40, and eluted by boiling 10 min in NuPage LDS sample buffer. Nonspecific mouse IgG (Jackson ImmunoResearch Laboratories) was used as the negative control.

Proteins (8% input lysate; 50% immunoprecipitate) were resolved on 10% SDS-PAGE gels (BioRad) and transferred to PVDF membranes (Immobilon-FL, Millipore). Blots were blocked and incubated overnight with primary antibodies against *O*-GlcNAc (mouse monoclonal antibody CTD_110.6_; Covance MMS-248R, diluted 1:5,000), OGT (AL25, diluted 1:2,000), lamins A/C (5G4; diluted 1:2,500) or lamin A (L1293; Sigma-Aldrich; diluted 1:2,000). Secondary HRP-conjugated anti-mouse (115-035-174) and anti-rabbit (211-032-171) IgG antibodies (both from Jackson ImmunoResearch Laboratories) were used at 1:10,000 dilution. Secondary anti-mouse IgM antibody (A8786; Sigma) was used at 1:5,000 dilution. Blots were developed using ECL (Pierce), and quantified from five independent experiments using ImageJ software.

### Succinylated wheat germ agglutinin (sWGA) pulldown

Mice liver nuclear lysates (100 ug per sample) were incubated 1 h (4°C) with protein A/G agarose beads (sc-2003; Santa Cruz Biotechnology), then centrifuged. The resulting supernatants (cleared extracts) were transferred to new tubes and incubated overnight (rotating; 4°C) with 40 ul succinylated wheat germ agglutinin (sWGA)-agarose (Vector Lab). After washing four times with PBS containing 0.2% NP-40, proteins were eluted from the beads by boiling for 10 min in NuPage LDS sample buffer, and resolved by SDS-PAGE. The captured proteins were analyzed by immunoblotting as described above.

### Mice

The male mice (mixed genetic background based on C57BL/6J and 129/Sv strains, backcrossed with C57BL/6J for at least six generations) were housed in a temperature-controlled (22°C) facility with a strict 12 h light/dark cycle. Mice had free access to food and water at all times. STZ (S0130; Sigma-Aldrich) was prepared in a sodium citrate buffer (50 mmol/l, pH 4.5) immediately before injections. STZ-treated mice were treated with two intraperitoneal injections of STZ (100 mg/kg) with a 1-day interval. Seven days after the first STZ injection, mice were included in a fasting-refeeding experiment. Mice were either fasted for 24 h, or fasted for 24 h and refed for 12 h on a diet containing 64% carbohydrates, 31.5% protein, 4.5% fat, and no cholesterol (SDS RM no.1 maintenance, Special Diets Services, UK). The mice were euthanized by cervical dislocation at 8:00 AM and liver tissues were snap-frozen in liquid nitrogen and stored at −80°C until further analysis. All use of animals was approved and registered by the Norwegian Animal Research authority.

### Peptide arrays and antibody epitope mapping

To generate peptide arrays, human lamin A (P02545, full length), human lamin C (NP_005563, full length), mouse lamin A (P48678, 588-end) and rat lamin A (P48679, 588-end) were synthesized as 20-mer peptides with three amino acid offsets on cellulose membranes using a Multipep automated peptide synthesizer (INTAVIS Bioanalytical Instruments AG, Koeln, Germany) as described (88). Peptide array membranes were blocked 2 h in 1% casein in TBST (Tris-buffered saline with 1% tween) at 22-25°C, then incubated overnight at 4°C with primary antibody (5G4 at 1:2,500 dilution; L1283 at 1:2,000) in TBST/1% casein. Membranes were then washed three times in TBST (10 min each) and incubated with affinity-purified horseradish-peroxidase-conjugated polyclonal anti-mouse IgG HRP (NA931V) or anti-rabbit IgG HRP (NA934V), both from GE HealthCare. Blots were developed using ECL Prime (RPN 2232, GE HealthCare) and chemiluminescence signals were detected using Las 1000 (Fujifilm, Tokyo, Japan). DNAStar Lasergene software (Madison, Wisconsin) was used to generate peptide sequence alignments.

### Purification of recombinant lamin tails

Recombinant His-and-T7-tagged lamin tail polypeptides were expressed in *E. coli* BL-21, purified using nickel NTA-agarose and stored in buffer (50 mM NaHPO_4_ pH 8.0, 300 mM NaCl, 100 mM imidazole, 0.5 mM PMSF) at −80°C until use, as described (80).

### In vitro *O*-GlcNAcylation reactions

Each reaction contained 1 ug recombinant lamin tails, 1 Unit CIP (New England Biolabs) and 10 mM UDP-GlcNAc (Sigma), plus or minus 1 ug purified recombinant active His-tagged OGT enzyme in a final reaction volume of 20 ul in 50 mM Tris-HCl pH 7.4. Reactions were incubated 2 h at 22-25°C, then overnight at 4°C. Reactions were stopped by adding 4X SDS sample buffer and 33% of each reaction was resolved on 4-12% Bis-Tris NuPage gels (Invitrogen), then transferred to nitrocellulose membranes, blocked 1 h at 22-25°C with 3% BSA in PBS, and incubated overnight at 4°C with affinity-purified *O*-GlcNAc-specific mouse antibody CTD_110.6_ (diluted 1:1000) or with a mixture of CTD_110.6_ antibody and competing (free) GlcNAc sugar (100 mM). Secondary antibodies were horseradish peroxidase-coupled anti-mouse IgM (Santa Cruz SC-2064; diluted 1:10,000) and anti-T7-tag (Novagen 69048-3; diluted 1:100,000).

### Mass Spectrometry

Recombinant purified His-tagged lamin A tails (wildtype, S612A/T643A, Δ35), *O*-GlcNAcylated in vitro as described above, were reduced for 1 h with dithiothreitol (DTT, Sigma) at a molar ratio of 20:1 (DTT:cysteine), carbamidomethylated for 1 h in the dark with iodoacetamide (IAA, Sigma) at a molar ratio of 3:1 (IAA:DTT), then proteolytically digested in 100 mM ammonium bicarbonate at a molar ratio of 1:20 (enzyme: substrate) by AspN (Roche, Inc), quenched with glacial acetic acid (to pH 3-4), and stored at −35°C. An additional chymotrypsin (Roche) digest was performed using similar conditions. For mass spectrometric analysis a fraction of the digest was pressure loaded onto a precolumn (360 μm o.d. × 75 μm i.d., fused silica capillary) packed with 6-8 cm of C18 reverse-phase resin (5-20 μm irregular diameter, 120 Á pore size, YMC). After a desalting rinse with 0.1 M acetic acid, the precolumn was connected via polytetrafluoroethylene tubing (0.06 in. o.d. × 0.012 in. o.d., Zeus Industrial Products) to the end of an analytical column (360 μm o.d. × 50 μm i.d. fused silica capillary) packed 6-8 cm with C18 reverse-phase resin (5 μm diameter, 120 Á pore size, YMC) and equipped with an electrospray emitter tip (89). Peptides were gradient eluted directly into the mass spectrometer with an Agilent 1100 series binary LC pump at a flow rate of ~60 nl/min with the following gradient: 0-60% B in 60 min, 60-100% B in 65 min, hold at 100% B for 70 min (solvent A: 0.1 M acetic acid, solvent B: 70% acetonitrile). Mass spectra were acquired with a modified front-end electron transfer-enabled (90) high-resolution LTQ-FT or LTQ-Orbitrap mass spectrometer (Thermo Scientific). Mass analyses were completed with one high resolution MS1 (60,000 at m/z 400) scan followed by 6 CAD and ETD MS2 scans acquired with the LTQ operating in either data-dependent or targeting mode. Azulene was used for ETD reactions with times of 30-50 ms.

Data from MS/MS analyses were searched against human lamin A using the Open Mass Spectrometry Search Algorithm (OMSSA; (91). OMSSA search tolerances were ± 0.01 Da and ± 0.35 Da for precursor and product ion masses, respectively. For ETD spectra, search parameters were set to exclude reduced charge species from peak lists prior to searching. Database searches were completed using either AspN or no enzyme specifications and allowing up to 3 missed cleavages. Specified variable modifications: carbamidomethylation of Cys, oxidation of Met, and *O*-GlcNAcylation of Ser and Thr. While OMSSA searches were used as a guide, all spectra and *O*-GlcNAc site modifications were validated by manual interpretation of the raw data.

## ACKNOWLEDGMENTS

We gratefully acknowledge funding from the Progeria Research Foundation (#2013-47 to KLW), Johns Hopkins University Claude D. Pepper Older Americans Independence Center National Institutes of Health NIA P30AG021334 (KLW), US-Israel Binational Science Foundation (#2011270 to YG and KLW), National Institutes of Health T32GM007445 (DS), National Institutes of Health GM037537 (DFH), Research Council of Norway (CRC, QF, LMGW), and the University of Oslo and Anders Jahre Foundation (QF, LMGW). We thank Jason Berk, Cliff Jenkins-Houk, and Joana Eid for early contributions to this project, and Natasha Zachara, Gerald Hart, and NHLBI Core C4 (Johns Hopkins School of Medicine) for purified OGT enzyme and *O*-GlcNAc antibodies.

## AUTHOR CONTRIBUTIONS

DNS and KLW conceived and designed the study, analyzed results, and drafted the paper, with supporting analysis and figure preparation by QF, LMGW, AW, TD, and YG. QF, LMGW, CRC, AF, and SBP designed, performed, and analyzed supporting and/or final experiments for Figure 1. CRC designed and performed the experiments in Supplemental Figure 1. DNS and YG performed and analyzed the experiments in Figure 2. DNS prepared samples for Figure 3. AW, JS, and DFH designed, performed, and analyzed the experiments in Figures 3-5. All authors contributed intellectually, reviewed results, and approved the final manuscript.

## REFERENCES

1. Simon, D. N., and Wilson, K. L. (2011) The nucleoskeleton as a genome-associated dynamic ‘network of networks’. Nat Rev Mol Cell Biol 12, 695–708.

2. Worman, H. J., and Schirmer, E. C. (2015) Nuclear membrane diversity: underlying tissue-specific pathologies in disease? Curr Opin Cell Biol 34, 101–112.

3. Gruenbaum, Y., and Foisner, R. (2015) Lamins: nuclear intermediate filament proteins with fundamental functions in nuclear mechanics and genome regulation. Annu Rev Biochem 84, 131–164.

4. Dittmer, T. A., and Misteli, T. (2011) The lamin protein family. Genome Biol 12, 222

5. Peter, A., and Stick, R. (2012) Evolution of the lamin protein family: What introns can tell. Nucleus 3

6. Koreny, L., and Field, M. C. (2016) Ancient Eukaryotic Origin and Evolutionary Plasticity of Nuclear Lamina. Genome Biol Evol 8, 2663–2671.

7. Burke, B., and Stewart, C. L. (2014) Functional architecture of the cell’s nucleus in development, aging, and disease. Current topics in developmental biology 109, 1–52.

8. Young, J., Morbois-Trabut, L., Couzinet, B., Lascols, O., Dion, E., Bereziat, V., Feve, B., Richard, I., Capeau, J., Chanson, P., and Vigouroux, C. (2005) Type A insulin resistance syndrome revealing a novel lamin A mutation. Diabetes 54, 1873–1878.

9. Guenantin, A. C., Briand, N., Bidault, G., Afonso, P., Bereziat, V., Vatier, C., Lascols, O., Caron-Debarle, M., Capeau, J., and Vigouroux, C. (2014) Nuclear envelope-related lipodystrophies. Semin Cell Dev Biol 29, 148–157.

10. Vigouroux, C., Magre, J., Vantyghem, M. C., Bourut, C., Lascols, O., Shackleton, S., Lloyd, D. J., Guerci, B., Padova, G., Valensi, P., Grimaldi, A., Piquemal, R., Touraine, P., Trembath, R. C., and Capeau, J. (2000) Lamin A/C gene: sex-determined expression of mutations in Dunnigan-type familial partial lipodystrophy and absence of coding mutations in congenital and acquired generalized lipoatrophy. Diabetes 49, 1958–1962.

11. Rizza, R. A. (2010) Pathogenesis of fasting and postprandial hyperglycemia in type 2 diabetes: implications for therapy. Diabetes 59, 2697–2707.

12. Dutour, A., Roll, P., Gaborit, B., Courrier, S., Alessi, M. C., Tregouet, D. A., Angelis, F., Robaglia-Schlupp, A., Lesavre, N., Cau, P., Levy, N., Badens, C., and Morange, P. E. (2011) High prevalence of laminopathies among patients with metabolic syndrome. Hum Mol Genet 20, 3779–3786.

13. Decaudain, A., Vantyghem, M. C., Guerci, B., Hecart, A. C., Auclair, M., Reznik, Y., Narbonne, H., Ducluzeau, P. H., Donadille, B., Lebbe, C., Bereziat, V., Capeau, J., Lascols, O., and Vigouroux, C. (2007) New metabolic phenotypes in laminopathies: LMNA mutations in patients with severe metabolic syndrome. J Clin Endocrinol Metab 92, 4835–4844.

14. Merideth, M. A., Gordon, L. B., Clauss, S., Sachdev, V., Smith, A. C., Perry, M. B., Brewer, C. C., Zalewski, C., Kim, H. J., Solomon, B., Brooks, B. P., Gerber, L. H., Turner, M. L., Domingo, D. L., Hart, T. C., Graf, J., Reynolds, J. C., Gropman, A., Yanovski, J. A., Gerhard-Herman, M., Collins, F. S., Nabel, E. G., Cannon, R. O., 3rd, Gahl, W. A., and Introne, W. J. (2008) Phenotype and course of Hutchinson-Gilford progeria syndrome. N Engl J Med 358, 592–604.

15. Eriksson, M., Brown, W. T., Gordon, L. B., Glynn, M. W., Singer, J., Scott, L., Erdos, M. R., Robbins, C. M., Moses, T. Y., Berglund, P., Dutra, A., Pak, E., Durkin, S., Csoka, A. B., Boehnke, M., Glover, T. W., and Collins, F. S. (2003) Recurrent de novo point mutations in lamin A cause Hutchinson-Gilford progeria syndrome. Nature 423, 293–298.

16. Fukuchi, K., Katsuya, T., Sugimoto, K., Kuremura, M., Kim, H. D., Li, L., and Ogihara, T. (2004) LMNA mutation in a 45 year old Japanese subject with Hutchinson-Gilford progeria syndrome. J Med Genet 41, e67

17. Vidak, S., and Foisner, R. (2016) Molecular insights into the premature aging disease progeria. Histochem Cell Biol 145, 401–417.

18. Gordon, L. B., Kleinman, M. E., Miller, D. T., Neuberg, D. S., Giobbie-Hurder, A., Gerhard-Herman, M., Smoot, L. B., Gordon, C. M., Cleveland, R., Snyder, B. D., Fligor, B., Bishop, W. R., Statkevich, P., Regen, A., Sonis, A., Riley, S., Ploski, C., Correia, A., Quinn, N., Ullrich, N. J., Nazarian, A., Liang, M. G., Huh, S. Y., Schwartzman, A., and Kieran, M. W. (2012) Clinical trial of a farnesyltransferase inhibitor in children with Hutchinson-Gilford progeria syndrome. Proc Natl Acad Sci U S A 109, 16666–16671.

19. Wang, Z., Udeshi, N. D., Slawson, C., Compton, P. D., Sakabe, K., Cheung, W. D., Shabanowitz, J., Hunt, D. F., and Hart, G. W. (2010) Extensive crosstalk between O-GlcNAcylation and phosphorylation regulates cytokinesis. Sci Signal 3, ra2

20. Hart, G. W., and Copeland, R. J. (2010) Glycomics hits the big time. Cell 143, 672–676.

21. Shafi, R., Iyer, S. P., Ellies, L. G., O’Donnell, N., Marek, K. W., Chui, D., Hart, G. W., and Marth, J. D. (2000) The O-GlcNAc transferase gene resides on the X chromosome and is essential for embryonic stem cell viability and mouse ontogeny. Proc Natl Acad Sci U S A 97, 5735–5739.

22. Hart, G. W. (2014) Minireview series on the thirtieth anniversary of research on O-GlcNAcylation of nuclear and cytoplasmic proteins: Nutrient regulation of cellular metabolism and physiology by O-GlcNAcylation. J Biol Chem 289, 34422–34423.

23. Levine, Z. G., and Walker, S. (2016) The Biochemistry of O-GlcNAc Transferase: Which Functions Make It Essential in Mammalian Cells? Annu Rev Biochem 85, 631–657.

24. Bond, M. R., and Hanover, J. A. (2013) O-GlcNAc cycling: a link between metabolism and chronic disease. Annu Rev Nutr 33, 205–229.

25. Copeland, R. J., Bullen, J. W., and Hart, G. W. (2008) Cross-talk between GlcNAcylation and phosphorylation: roles in insulin resistance and glucose toxicity. Am J Physiol Endocrinol Metab 295, E17–28.

26. Hardiville, S., and Hart, G. W. (2014) Nutrient regulation of signaling, transcription, and cell physiology by O-GlcNAcylation. Cell Metab 20, 208–213.

27. Jensen, R. V., Zachara, N. E., Nielsen, P. H., Kimose, H. H., Kristiansen, S. B., and Botker, H. E. (2013) Impact of O-GlcNAc on cardioprotection by remote ischaemic preconditioning in non-diabetic and diabetic patients. Cardiovasc Res 97, 369–378.

28. Ramirez-Correa, G. A., Jin, W., Wang, Z., Zhong, X., Gao, W. D., Dias, W. B., Vecoli, C., Hart, G. W., and Murphy, A. M. (2008) O-linked GlcNAc modification of cardiac myofilament proteins: a novel regulator of myocardial contractile function. Circ Res 103, 1354–1358.

29. Zhang, K., Yin, R., and Yang, X. (2014) O-GlcNAc: A Bittersweet Switch in Liver. Front Endocrinol (Lausanne) 5, 221

30. Like, A. A., and Rossini, A. A. (1976) Streptozotocin-induced pancreatic insulitis: new model of diabetes mellitus. Science 193, 415–417.

31. Simon, D. N., and Wilson, K. L. (2013) Partners and post-translational modifications of nuclear lamins. Chromosoma 122, 13–31.

32. Kochin, V., Shimi, T., Torvaldson, E., Adam, S. A., Goldman, A., Pack, C. G., Melo-Cardenas, J., Imanishi, S. Y., Goldman, R. D., and Eriksson, J. E. (2014) Interphase phosphorylation of lamin A. J Cell Sci 127, 2683–2696.

33. Navarro, C. L., De Sandre-Giovannoli, A., Bernard, R., Boccaccio, I., Boyer, A., Genevieve, D., Hadj-Rabia, S., Gaudy-Marqueste, C., Smitt, H. S., Vabres, P., Faivre, L., Verloes, A., Van Essen, T., Flori, E., Hennekam, R., Beemer, F. A., Laurent, N., Le Merrer, M., Cau, P., and Levy, N. (2004) Lamin A and ZMPSTE24 (FACE-1) defects cause nuclear disorganization and identify restrictive dermopathy as a lethal neonatal laminopathy. Hum Mol Genet 13, 2493–2503.

34. Ferraro, A., Eufemi, M., Cervoni, L., Marinetti, R., and Turano, C. (1989) Glycosylated forms of nuclear lamins. FEBS Lett 257, 241–246.

35. Alfaro, J. F., Gong, C. X., Monroe, M. E., Aldrich, J. T., Clauss, T. R., Purvine, S. O., Wang, Z., Camp, D. G., 2nd, Shabanowitz, J., Stanley, P., Hart, G. W., Hunt, D. F., Yang, F., and Smith, R. D. (2012) Tandem mass spectrometry identifies many mouse brain O-GlcNAcylated proteins including EGF domain-specific O-GlcNAc transferase targets. Proc Natl Acad Sci U S A 109, 7280–7285.

36. Pathak, S., Alonso, J., Schimpl, M., Rafie, K., Blair, D. E., Borodkin, V. S., Schuttelkopf, A. W., Albarbarawi, O., and van Aalten, D. M. (2015) The active site of O-GlcNAc transferase imposes constraints on substrate sequence. Nat Struct Mol Biol 22, 744–750.

37. Herrmann, H., Kreplak, L., and Aebi, U. (2004) Isolation, characterization, and in vitro assembly of intermediate filaments. Methods Cell Biol 78, 3–24.

38. Zhao, P., Schulz, T. C., Sherrer, E. S., Weatherly, D. B., Robins, A. J., and Wells, L. (2015) The human embryonic stem cell proteome revealed by multidimensional fractionation followed by tandem mass spectrometry. Proteomics 15, 554–566.

39. Jochmann, R., Holz, P., Sticht, H., and Sturzl, M. (2014) Validation of the reliability of computational O-GlcNAc prediction. Biochim Biophys Acta 1844, 416–421.

40. Schutkowski, M., Reimer, U., Panse, S., Dong, L., Lizcano, J. M., Alessi, D. R., and Schneider-Mergener, J. (2004) High-content peptide microarrays for deciphering kinase specificity and biology. Angew Chem Int Ed Engl 43, 2671–2674.

41. Liu, X., Li, L., Wang, Y., Yan, H., Ma, X., Wang, P. G., and Zhang, L. (2014) A peptide panel investigation reveals the acceptor specificity of O-GlcNAc transferase. FASEB J 28, 3362–3372.

42. Kao, H. J., Huang, C. H., Bretana, N. A., Lu, C. T., Huang, K. Y., Weng, S. L., and Lee, T. Y. (2015) A two-layered machine learning method to identify protein O-GlcNAcylation sites with O-GlcNAc transferase substrate motifs. BMC Bioinformatics 16 Suppl 18, S10

43. Gupta, R., and Brunak, S. (2002) Prediction of glycosylation across the human proteome and the correlation to protein function. Pac Symp Biocomput, 310–322.

44. Jia, C. Z., Liu, T., and Wang, Z. P. (2013) O-GlcNAcPRED: a sensitive predictor to capture protein O-GlcNAcylation sites. Mol Biosyst 9, 2909–2913.

45. Berk, J. M., Maitra, S., Dawdy, A. W., Shabanowitz, J., Hunt, D. F., and Wilson, K. L. (2013) O-Linked beta-N-acetylglucosamine (O-GlcNAc) regulates emerin binding to barrier to autointegration factor (BAF) in a chromatin-and lamin B-enriched “niche”. J Biol Chem 288, 30192–30209.

46. Barrowman, J., Hamblet, C., Kane, M. S., and Michaelis, S. (2012) Requirements for efficient proteolytic cleavage of prelamin A by ZMPSTE24. PLoS One 7, e32120

47. Casasola, A., Scalzo, D., Nandakumar, V., Halow, J., Recillas-Targa, F., Groudine, M., and Rincon-Arano, H. (2016) Prelamin A processing, accumulation and distribution in normal cells and laminopathy disorders. Nucleus 7, 84–102.

48. He, M., Wang, Y., and Li, W. (2009) PPI finder: a mining tool for human protein-protein interactions. PLoS One 4, e4554

49. Shimi, T., Kittisopikul, M., Tran, J., Goldman, A. E., Adam, S. A., Zheng, Y., Jaqaman, K., and Goldman, R. D. (2015) Structural organization of nuclear lamins A, C, B1, and B2 revealed by superresolution microscopy. Mol Biol Cell 26, 4075–4086.

50. Xie, W., Chojnowski, A., Boudier, T., Lim, J. S., Ahmed, S., Ser, Z., Stewart, C., and Burke, B. (2016) A-type Lamins Form Distinct Filamentous Networks with Differential Nuclear Pore Complex Associations. Curr Biol 26, 2651–2658.

51. Jung, H. J., Coffinier, C., Choe, Y., Beigneux, A. P., Davies, B. S., Yang, S. H., Barnes, R. H., 2nd, Hong, J., Sun, T., Pleasure, S. J., Young, S. G., and Fong, L. G. (2012) Regulation of prelamin A but not lamin C by miR-9, a brain-specific microRNA. Proc Natl Acad Sci U S A 109, E423–431.

52. Lopez-Mejia, I. C., de Toledo, M., Chavey, C., Lapasset, L., Cavelier, P., Lopez-Herrera, C., Chebli, K., Fort, P., Beranger, G., Fajas, L., Amri, E. Z., Casas, F., and Tazi, J. (2014) Antagonistic functions of LMNA isoforms in energy expenditure and lifespan. EMBO Rep 15, 529–539.

53. Coffinier, C., Jung, H. J., Li, Z., Nobumori, C., Yun, U. J., Farber, E. A., Davies, B. S., Weinstein, M. M., Yang, S. H., Lammerding, J., Farahani, J. N., Bentolila, L. A., Fong, L. G., and Young, S. G. (2010) Direct synthesis of lamin A, bypassing prelamin a processing, causes misshapen nuclei in fibroblasts but no detectable pathology in mice. J Biol Chem 285, 20818–20826.

54. Evangelisti, C., Cenni, V., and Lattanzi, G. (2016) Potential therapeutic effects of the MTOR inhibitors for preventing ageing and progeria-related disorders. Br J Clin Pharmacol 82, 1229–1244.

55. Mendelsohn, A. R., and Larrick, J. W. (2011) Rapamycin as an antiaging therapeutic?: targeting mammalian target of rapamycin to treat Hutchinson-Gilford progeria and neurodegenerative diseases. Rejuvenation Res 14, 437–441.

56. Shimobayashi, M., and Hall, M. N. (2014) Making new contacts: the mTOR network in metabolism and signalling crosstalk. Nat Rev Mol Cell Biol 15, 155–162.

57. Park, Y. E., Hayashi, Y. K., Bonne, G., Arimura, T., Noguchi, S., Nonaka, I., and Nishino, I. (2009) Autophagic degradation of nuclear components in mammalian cells. Autophagy 5, 795–804.

58. Ramos, F. J., Chen, S. C., Garelick, M. G., Dai, D. F., Liao, C. Y., Schreiber, K. H., MacKay, V. L., An, E. H., Strong, R., Ladiges, W. C., Rabinovitch, P. S., Kaeberlein, M., and Kennedy, B. K. (2012) Rapamycin reverses elevated mTORC1 signaling in lamin A/C-deficient mice, rescues cardiac and skeletal muscle function, and extends survival. Sci Transl Med 4, 144ra103

59. Choi, J. C., Muchir, A., Wu, W., Iwata, S., Homma, S., Morrow, J. P., and Worman, H. J. (2012) Temsirolimus activates autophagy and ameliorates cardiomyopathy caused by lamin A/C gene mutation. Sci Transl Med 4, 144ra102

60. Cattin, M. E., Wang, J., Weldrick, J. J., Roeske, C. L., Mak, E., Thorn, S. L., DaSilva, J. N., Wang, Y., Lusis, A. J., and Burgon, P. G. (2015) Deletion of MLIP (muscle-enriched A-type lamin-interacting protein) leads to cardiac hyperactivation of Akt/mammalian target of rapamycin (mTOR) and impaired cardiac adaptation. J Biol Chem 290, 26699–26714.

61. Choi, J. C., and Worman, H. J. (2013) Reactivation of autophagy ameliorates LMNA cardiomyopathy. Autophagy 9, 110–111.

62. Ramos, F. J., Kaeberlein, M., and Kennedy, B. K. (2013) Elevated MTORC1 signaling and impaired autophagy. Autophagy 9, 108–109.

63. Ullrich, N. J., Kieran, M. W., Miller, D. T., Gordon, L. B., Cho, Y. J., Silvera, V. M., Giobbie-Hurder, A., Neuberg, D., and Kleinman, M. E. (2013) Neurologic features of Hutchinson-Gilford progeria syndrome after lonafarnib treatment. Neurology 81, 427–430.

64. Florwick, A., Dharmaraj, T., Jurgens, J., Valle, D., and Wilson, K. L. (2017) LMNA Sequences of 60,706 Unrelated Individuals Reveal 132 Novel Missense Variants in A-Type Lamins and Suggest a Link between Variant p.G602S and Type 2 Diabetes. Front Genet 8, 79

65. Rankin, J., Auer-Grumbach, M., Bagg, W., Colclough, K., Nguyen, T. D., Fenton-May, J., Hattersley, A., Hudson, J., Jardine, P., Josifova, D., Longman, C., McWilliam, R., Owen, K., Walker, M., Wehnert, M., and Ellard, S. (2008) Extreme phenotypic diversity and nonpenetrance in families with the LMNA gene mutation R644C. Am J Med Genet A 146A, 1530–1542.

66. Harwood, K. R., and Hanover, J. A. (2014) Nutrient-driven O-GlcNAc cycling – think globally but act locally. J Cell Sci 127, 1857–1867.

67. de Leeuw, R., Gruenbaum, Y., and Medalia, O. (2017) Nuclear Lamins: Thin Filaments with Major Functions. Trends Cell Biol

68. Samwer, M., Schneider, M. W. G., Hoefler, R., Schmalhorst, P. S., Jude, J. G., Zuber, J., and Gerlich, D. W. (2017) DNA Cross-Bridging Shapes a Single Nucleus from a Set of Mitotic Chromosomes. Cell 170, 956–972 e923

69. Naetar, N., Ferraioli, S., and Foisner, R. (2017) Lamins in the nuclear interior - life outside the lamina. J Cell Sci 130, 2087–2096.

70. Liu, J., Lee, K. K., Segura-Totten, M., Neufeld, E., Wilson, K. L., and Gruenbaum, Y. (2003) MAN1 and emerin have overlapping function(s) essential for chromosome segregation and cell division in Caenorhabditis elegans. Proc Natl Acad Sci U S A 100, 4598–4603.

71. Liu, J., Rolef Ben-Shahar, T., Riemer, D., Treinin, M., Spann, P., Weber, K., Fire, A., and Gruenbaum, Y. (2000) Essential roles for Caenorhabditis elegans lamin gene in nuclear organization, cell cycle progression, and spatial organization of nuclear pore complexes. Mol Biol Cell 11, 3937–3947.

72. Margalit, A., Segura-Totten, M., Gruenbaum, Y., and Wilson, K. L. (2005) Barrier-to-autointegration factor is required to segregate and enclose chromosomes within the nuclear envelope and assemble the nuclear lamina. Proc Natl Acad Sci U S A 102, 3290–3295.

73. Snider, N. T., and Omary, M. B. (2014) Post-translational modifications of intermediate filament proteins: mechanisms and functions. Nat Rev Mol Cell Biol 15, 163–177.

74. Roque, A., Ponte, I., and Suau, P. (2017) Post-translational modifications of the intrinsically disordered terminal domains of histone H1: effects on secondary structure and chromatin dynamics. Chromosoma 126, 83–91.

75. Uversky, V. N. (2017) Intrinsic disorder here, there, and everywhere, and nowhere to escape from it. Cell Mol Life Sci 74, 3065–3067.

76. Crisp, M., Liu, Q., Roux, K., Rattner, J. B., Shanahan, C., Burke, B., Stahl, P. D., and Hodzic, D. (2006) Coupling of the nucleus and cytoplasm: role of the LINC complex. J Cell Biol 172, 41–53.

77. Dittmer, T. A., Sahni, N., Kubben, N., Hill, D. E., Vidal, M., Burgess, R. C., Roukos, V., and Misteli, T. (2014) Systematic identification of pathological lamin A interactors. Mol Biol Cell 25, 1493–1510.

78. Tao, Y., Fang, P., Kim, S., Guo, M., Young, N. L., and Marshall, A. G. (2017) Mapping the contact surfaces in the Lamin A:AIMP3 complex by hydrogen/deuterium exchange FT-ICR mass spectrometry. PLoS One 12, e0181869

79. Oh, Y. S., Kim, D. G., Kim, G., Choi, E. C., Kennedy, B. K., Suh, Y., Park, B. J., and Kim, S. (2010) Downregulation of lamin A by tumor suppressor AIMP3/p18 leads to a progeroid phenotype in mice. Aging Cell 9, 810–822.

80. Simon, D. N., Zastrow, M. S., and Wilson, K. L. (2010) Direct actin binding to A-and B-type lamin tails and actin filament bundling by the lamin A tail. Nucleus 1, 264–272.

81. Xie, W., and Burke, B. (2017) Nuclear networking. Nucleus 8, 323–330.

82. Bond, M. R., and Hanover, J. A. (2015) A little sugar goes a long way: the cell biology of O-GlcNAc. J Cell Biol 208, 869–880.

83. Ma, J., and Hart, G. W. (2013) Protein O-GlcNAcylation in diabetes and diabetic complications. Expert Rev Proteomics 10, 365–380.

84. Ronningen, T., Shah, A., Oldenburg, A. R., Vekterud, K., Delbarre, E., Moskaug, J. O., and Collas, P. (2015) Prepatterning of differentiation-driven nuclear lamin A/C-associated chromatin domains by GlcNAcylated histone H2B. Genome Res 25, 1825–1835.

85. Bronshtein, I., Kepten, E., Kanter, I., Berezin, S., Lindner, M., Redwood, A. B., Mai, S., Gonzalo, S., Foisner, R., Shav-Tal, Y., and Garini, Y. (2015) Loss of lamin A function increases chromatin dynamics in the nuclear interior. Nat Commun 6, 8044

86. Cesarini, E., Mozzetta, C., Marullo, F., Gregoretti, F., Gargiulo, A., Columbaro, M., Cortesi, A., Antonelli, L., Di Pelino, S., Squarzoni, S., Palacios, D., Zippo, A., Bodega, B., Oliva, G., and Lanzuolo, C. (2015) Lamin A/C sustains PcG protein architecture, maintaining transcriptional repression at target genes. J Cell Biol 211, 533–551.

87. Perovanovic, J., Dell’Orso, S., Gnochi, V. F., Jaiswal, J. K., Sartorelli, V., Vigouroux, C., Mamchaoui, K., Mouly, V., Bonne, G., and Hoffman, E. P. (2016) Laminopathies disrupt epigenomic developmental programs and cell fate. Sci Transl Med 8, 335ra358

88. Frank, R., and Overwin, H. (1996) SPOT synthesis. Epitope analysis with arrays of synthetic peptides prepared on cellulose membranes. Methods Mol Biol 66, 149–169.

89. Udeshi, N. D., Compton, P. D., Shabanowitz, J., Hunt, D. F., and Rose, K. L. (2008) Methods for analyzing peptides and proteins on a chromatographic timescale by electron-transfer dissociation mass spectrometry. Nat Protoc 3, 1709–1717.

90. Earley, L., Anderson, L. C., Bai, D. L., Mullen, C., Syka, J. E., English, A. M., Dunyach, J. J., Stafford, G. C., Jr., Shabanowitz, J., Hunt, D. F., and Compton, P. D. (2013) Front-end electron transfer dissociation: a new ionization source. Anal Chem 85, 8385–8390.

91. Geer, L. Y., Markey, S. P., Kowalak, J. A., Wagner, L., Xu, M., Maynard, D. M., Yang, X., Shi, W., and Bryant, S. H. (2004) Open mass spectrometry search algorithm. J Proteome Res 3, 958–964.

92. Krimm, I., Ostlund, C., Gilquin, B., Couprie, J., Hossenlopp, P., Mornon, J. P., Bonne, G., Courvalin, J. C., Worman, H. J., and Zinn-Justin, S. (2002) The Ig-like structure of the C-terminal domain of lamin A/C, mutated in muscular dystrophies, cardiomyopathy, and partial lipodystrophy. Structure 10, 811–823.

